# Photoreceptor-induced LHL4 protects photosystem II in *Chlamydomonas reinhardtii*

**DOI:** 10.1101/2024.02.23.581703

**Authors:** Marie Dannay, Chloé Bertin, Eva Cavallari, Pascal Albanese, Dimitri Tolleter, Cécile Giustini, Mathilde Menneteau, Sabine Brugière, Yohann Couté, Giovanni Finazzi, Emilie Demarsy, Roman Ulm, Guillaume Allorent

**Author notes:** UMR9019 Genome Integrity & Cancers - CNRS, Université Paris-Saclay, Gustave Roussy Cancer Campus, 94800 Villejuif, France.

## Abstract

Photosynthesis, the fundamental process using light energy to convert CO_2_ to organic matter, is vital for life on Earth. It relies on capturing light through light-harvesting complexes in evolutionarily well-conserved photosystems (PS) I and II and on the conversion of light energy into chemical energy. Composition and organization of both photosystem core complexes are well conserved across evolution. PSII is particularly sensitive to photodamage but benefits from a large diversity of photoprotective mechanisms, finely tuned for the specific light conditions. Light Harvesting Complex protein family members (LHC and LHC-like families) have acquired a dual function during evolution. Members of the LHC antenna complexes of photosystems capture light energy whereas others dissipate excess energy that cannot be harnessed for photosynthesis. This process mainly occurs through non photochemical quenching (NPQ). In this work, we focus on the LHL4 protein, which is a LHC-like protein induced by UV-B and blue light photoreceptor signaling pathways in the model green microalgae *Chlamydomonas reinhardtii*. We demonstrate that alongside established NPQ effectors, LHL4 plays a key role in photoprotection, preventing singlet oxygen accumulation in PSII and promoting cell survival upon light stress. LHL4 protective function is distinct from that of NPQ-related proteins, as it specifically and uniquely binds to the transient monomeric form of the core PSII complex, safeguarding its integrity. LHL4 characterization expands our understanding of the interplay between light harvesting and photoprotection mechanisms upon light stress in photosynthetic microalgae.

## Introduction

Photosynthetic organisms encounter daily light fluctuations in their natural environments. As light is a source of energy for photosynthesis but can also be a source of photodamage, balancing light absorption and light dissipation is crucial for survival and fitness (1). Photosynthetic organisms have evolved different strategies to acclimate and respond to light fluctuations. These responses include complementary mechanisms that prevent, limit or repair photodamage and that are activated at different times upon perception of the light changes (2, 3). For example, dissipation of light energy through non photochemical quenching (NPQ) and scavenging of reactive oxygen species (ROS) are activated within minutes, playing crucial roles for survival under excessive light (4–7). Light-Harvesting Complex (LHC) proteins and the related LHC-like proteins have evolved different functions in light absorption and photoprotection from green algae to flowering plants (8). LHC-like family members contain one to four transmembrane domains each harbouring putative LHC motifs that may bind chlorophyll a/b, carotenoid, and lipids (8). In algae, including the green microalgae *Chlamydomonas reinhardtii* (abbreviated as Chlamydomonas hereafter), important NPQ effectors are the LHC-Like Stress Related (LHCSR) proteins LHCSR1 and LHCSR3 (9). In vascular plants, where LHCSR proteins are absent, Photosystem II Subunit S (PSBS) is a major player of NPQ (6). PSBS also contributes to NPQ in green microalgae, although to a lesser extent than the LHCSR proteins (10–13). Besides NPQ, plant photoprotection has been proposed to be modulated through the action of the LHC-like Early Light Induced Protein (ELIP) family. ELIPs possibly prevent the negative effect of free chlorophyll released during the turnover of photosystem II (PSII) (14). However, the exact function of other LHC-like protein remained unknown.

Photoprotective mechanisms are activated by photoreceptors that monitor the light environment (15). In green microalgae, expression of *LHCSR*s, *PSBS*, and *ELIP*s is highly light-inducible through blue light-activated phototropin (PHOT) and UV-B-activated UV Resistance Locus 8 (UVR8) photoreceptors (11, 16, 17). The early steps of UVR8 signaling are well conserved in the green lineage (18, 19), with UV-B-induced UVR8 monomerization and binding to Constitutively Photomorphogenic 1 (COP1), an E3 ubiquitin ligase playing a vital role in repressing light responses and photomorphogenesis in photosynthetic organisms (11, 20–24). UVR8 binding inhibits COP1, stabilizing transcription factors and thereby leading to transcriptional reprogramming (18, 25–27). In flowering plants, blue light gene expression relies on cryptochrome photoreceptor - COP1 mediated pathway (22, 26, 28), whereas phototropins seem to control blue light responses independently of early transcriptional changes (29, 30). In contrast, in Chlamydomonas, PHOT induces transcription of *LHCSR3* through a signaling cascade that involves COP1 (also known as LRS1/HIT1) (16, 27, 31, 32). Downstream of COP1, molecular players apparently diverged during evolution. The B-box transcription factor CONSTANS (CrCO) controls *LHCSR*s expression in response to both UV-B and high light (HL) in Chlamydomonas (27, 31), whereas the bZIP transcription factor HY5 plays the major role in the UV-B and blue light signaling pathway in flowering plants (33–37), but its role remains elusive in green algae (38).

Here, we identify the Chlamydomonas LHC-Like protein 4 (LHL4) as a crucial component of the photoprotection response. *LHL4* expression and LHL4 accumulation are strongly induced under UV-B and HL, in a UVR8- and PHOT-dependent manner, respectively. Induction involves transcriptional regulation through CrCO. Transcriptomic data corroborate the conclusion that CrCO plays a major role in UVR8-dependent transcriptional reprogramming. Finally, we suggest that LHL4-mediated photoprotection prevents ROS production during HL stress by directly interacting with the PSII monomer. The LHL4 protective function is prominent during a first phase of moderate HL stress, *i.e.* before activation of the canonical NPQ effectors, or when they do not provide sufficient protection under strong HL stress.

## Results and Discussion

### LHL4 is highly induced upon UV-B

We analysed the effect of UV-B on the membrane-enriched proteome of Chlamydomonas and identified several proteins potentially involved in photoprotection among those accumulating under 16 h UV-B (Fig. 1A). After UV-B exposure, 46 proteins showed a significant increase in abundance, and 9 proteins showed a significant decrease in abundance. Several proteins that showed increased accumulation in response to UV-B are involved in PSII biogenesis, stability, or repair (FTSH-like and DEG proteases (39), and members of the HCF family (40)), as well as in NPQ (the two LHC-like proteins LHCSR1 and LHCSR3) (Table S1). Interestingly, we identified a third LHC-like protein, namely LHL4, enriched more than 7 times under UV-B compared to control (Fig. 1A and Table S1). In a parallel RNA-Seq analysis, *LHL4* was also identified as highly induced at the transcript level in response to 1h exposure to UV-B, in a UVR8-dependent manner, along with 831 genes (Fig. 1B and Fig. S1 and Table S2). While LHCSRs proteins were detected in both experiments, PSBS was identified among the top-induced genes in our transcriptome dataset (Table S2), as expected (11, 24), but showed only limited accumulation at the protein level in the UV-B-treated samples (Table S1), consistent with its reported limited stability over time (10, 13). In total, 44 out of the 46 proteins accumulating in response to UV-B were also found to be transcriptionally induced (Fig. 1C and Table S3).

**Fig 1:**
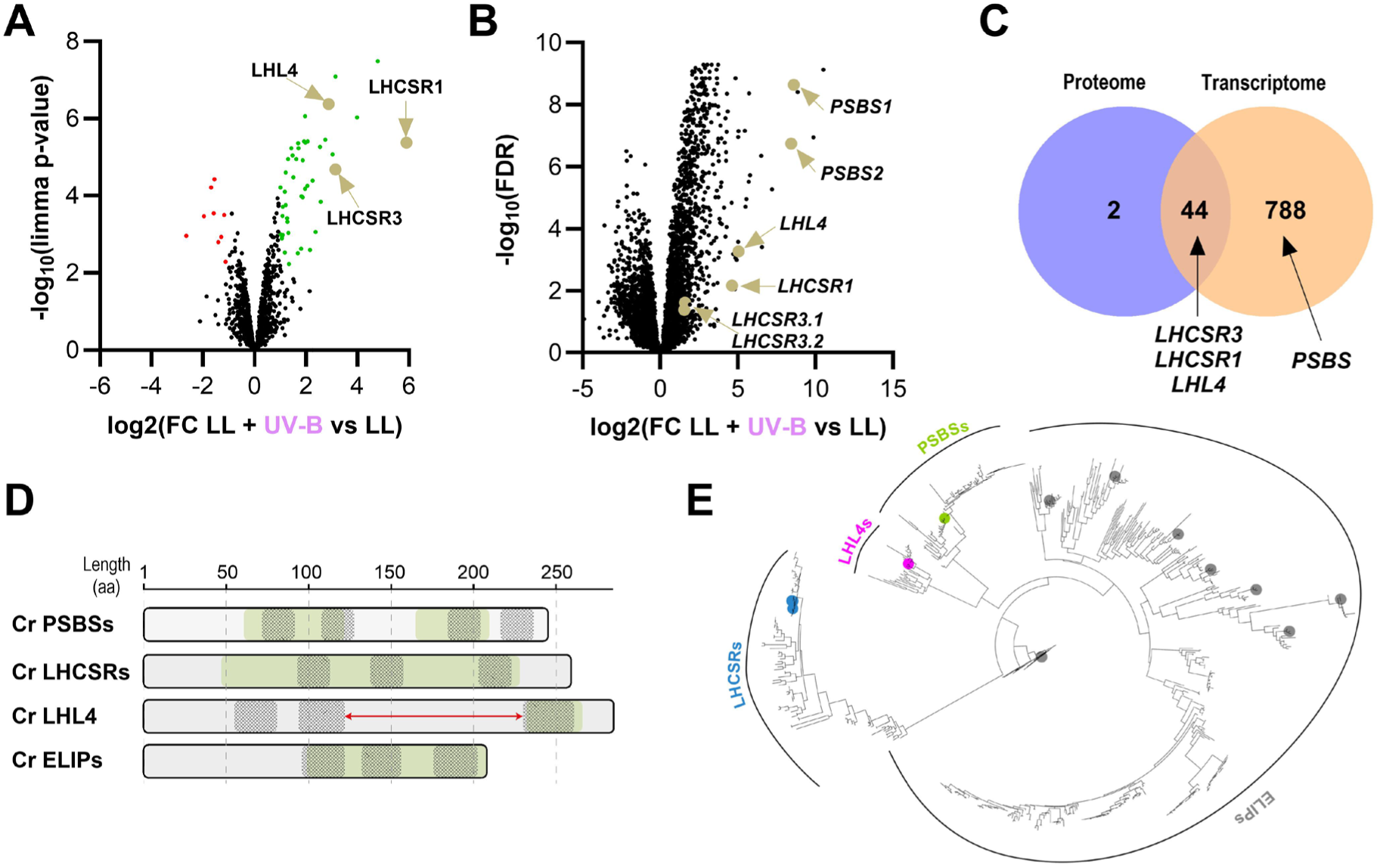
LHL4 is induced by UV-B in Chlamydomonas. **(A)** Mass spectrometry (MS)-based quantitative comparison of membrane-enriched proteomes from Chlamydomonas exposed to 16h of supplemental UV-B [Low Light (LL, 20 µmol photons m^-2^ s^-1^) + UV-B (0.2 mW cm*^-^*^2^)] compared to untreated control (LL). Volcano plot displaying the differential abundance of proteins in the membrane proteomes analysed by MS-based label-free quantitative proteomics. The volcano plot represents the -log10 (p-value), (limma p-value, y axis) plotted against the log_2_FC (LL + UV-B vs LL, x axis) for each quantified protein. Green and red dots represent proteins significantly enriched in LL + UV-B and in LL samples, respectively (log_2_FC ≥ 1 and -log_10_(p-value) ≥ 2.11, corresponding to a Benjamini-Hochberg FDR < 1 %). Dots representing LHL4, LHCSR1, and LHCSR3 proteins are indicated. **(B)** RNA-Seq analysis of Chlamydomonas exposed to 1h LL + UV-B compared to LL. Volcano plot displaying the differential abundance of transcripts by representing the -log_10_ (FDR), (y axis) plotted against the log_2_FC (LL + UV-B vs LL, x axis). Dots representing *LHL4*, *PSBS1*/*PSBS2* (encoding PSBS), *LHCSR1*, and *LHCSR3.1*/*LHCSR3.2* (encoding LHCSR3) transcripts are indicated. **(C)** Venn diagram showing the overlap of proteins (blue) and transcripts (orange) significantly enriched in LL + UV-B (Tables S1 and S2). **(D)** Scaled schematic representation of LHC-Like protein sizes and the locations of predicted chlorophyll a/b binding domains (green, predicted using the Superfamily database) and transmembrane domains (grey motifs) in LHL4 and in consensus Chlamydomonas ELIP, LHCSR, and PSBS like protein sequences. The red arrow indicates the presence of an exceptionally long loop in LHL4 compared to the other proteins. aa, amino acids. **(E)** Phylogenetic tree generated from aligned protein sequences of LHC-Like LHCSRs, LHL4s, PSBSs and ELIPs homologs. LHCSRs (blue), LHL4 (magenta), PSBSs (green), and ELIPs (grey) proteins from Chlamydomonas are approximatively emphasized by colored dots.

LHL4 is a 285-amino acid protein with three transmembrane domains that differs from the other LHC-Like proteins because it contains an exceptionally long predicted loop between the second and third transmembrane domains (Fig. 1D) (41, 42). LHL4 homologs are uniquely found in green microalgae (43, 44), where they are closely related to PSBS, but belong to a different clade (Fig. 1E).

### *LHL4* expression is controlled by UVR8 and PHOT photoreceptors, and depends on CrCOP1 and CrCO

*LHL4* gene expression was found to be rapidly and transiently induced in response to both UV-B and HL (Fig. 2A) (24, 41, 42). *LHL4* showed a higher induction by UV-B than HL at both transcript and protein levels. The LHL4 protein level rapidly increased up to 4 h and remained stable for 8 h upon both light treatments (Fig. 2B). However, the LHL4 abundance rapidly decreased when cells were returned to low light (LL), unlike LHCSR1 and LHCSR3, which remained stable for at least 8 h post-treatment (Fig. S2) (45).

**Fig 2:**
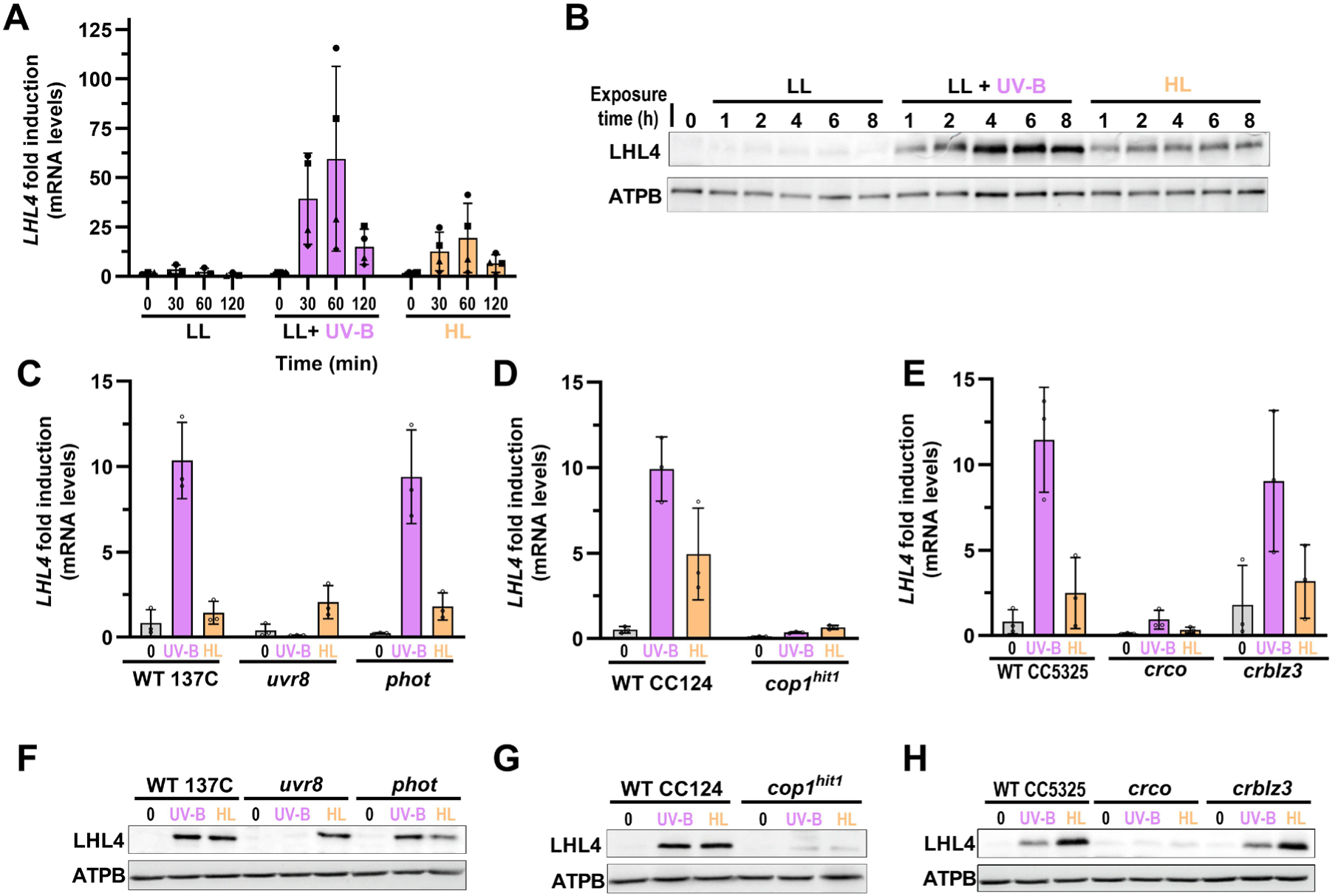
LHL4 expression is controlled by UVR8 and PHOT photoreceptor signaling pathways. **(A)** RT-qPCR analysis of *LHL4* expression in WT 137C cells grown under LL (20 µmol photons m^-2^ s^-1^) and then transferred to LL supplemented with UV-B (0.2 mW cm*^-^*^2^, LL + UV-B), or HL (300 µmol photons m^-2^ s^-1^) for the indicated times. The data were normalized to the *LHL4* levels at time 0 for each condition. Individual data points of biological replicates and means ± SD are shown (*n* = 4). **(B)** Immunodetection of LHL4 in cells exposed for up to 8 h under LL, LL + UV-B, or HL. ATP synthase beta subunit (ATPB) levels were used as loading control. **(C-E)** RT-qPCR analysis of *LHL4* expression in **(C)** *uvr8* and *phot*, **(D)** *cop1^hit1^*and **(E)** *crco* and *crblz3* cells grown under LL (0), or exposed for 1 h to LL + UV-B, or HL. Data are normalized to levels in respective WTs (strains 137C, CC124, CC5325) under LL (“0”). Values of independent measurements and means ± SD are shown (*n* = 3). **(F-H)** Immunodetection of LHL4 in **(F)** *uvr8* and *phot*, **(G)** *cop1^hit1^*, **(H)** *crco* and *crblz3*, as well as their respective WT cells grown under LL (0), or 6 h LL + UV-B, or HL. ATPB was used as loading control.

As *LHL4* expression is induced in response to UV-B and HL, we examined the accumulation of *LHL4* transcripts and LHL4 proteins in *uvr8* and *phot* mutants. UV-B-dependent induction of *LHL4* expression was abolished in the *uvr8* mutant, but not affected in a *phot* mutant (Fig. 2C). Consistently, UV-B-induced LHL4 protein accumulation was absent in *uvr8* but comparable to wild type (WT) in *phot* cells (Fig. 2F). Under HL, a comparably weak *LHL4* induction was observed in both photoreceptor mutants and WT cells (Fig. 2C). However, accumulation of LHL4 protein was reduced in response to HL in the *phot* mutant compared to WT and *uvr8* (Fig. 2F), similar to LHCSR3 (Fig. S3A) (16). Altogether, our data demonstrate that UV-B- and HL-induced LHL4 accumulation is mediated by UVR8 and, at least partially, by PHOT, respectively. We thus examined the involvement of downstream signaling components on *LHL4* induction and LHL4 accumulation. We first used the *cop1^hit1^* loss-of-function mutant strain that contains an Arg-1256-to-Pro mutation (CrCOP1^R1256P^) in the C-terminal WD40 domain (11, 32). Under both UV-B and HL, the *cop1^hit1^* mutant exhibited a much weaker induction of *LHL4* transcript and reduced LHL4 protein accumulation in comparison to WT (Fig. 2D and 2G). This result is similar to the reduced accumulation of LHCSR1 and LHCSR3 (Fig. S3B), as well as PSBS (11), confirming that the UV-B and HL signaling pathways converge at the level of COP1. It is also of note that *cop1^hit1^* is more strongly affected in HL-induced LHCSR3 and LHCSR1 accumulation than *phot* (Fig. S3A and S3B), suggesting that additional HL induced signaling pathway(s) converge at COP1.

We next investigated transcription factors potentially involved in *LHL4* regulation upon UV-B and HL by comparing LHL4 levels in *crco* (27, 31) and *crblz3* (CrBLZ3 is a putative AtHY5 ortholog, (38)) mutants with WT. At the transcriptome-wide level, we found that both CrCO and CrBLZ3 are required for UV-B regulated gene expression with CrCO playing a major role (Fig. S4A, B, and Table S2). In agreement, accumulation of *LHL4* mRNA and LHL4 protein in response to both UV-B and HL was severely impaired in *crco*, but was similar between *crblz3* and WT (Fig. 2E, H, and Table S2). We conclude that UV-B- and HL-dependent accumulation of LHL4 depends on UVR8 and partially on PHOT, respectively, in both cases involving CrCO-dependent transcriptional activation of *LHL4* expression.

### LHL4 prevents PSII photoinhibition upon exposure to HL

The sequence homology of LHL4 with LHCSR and PSBS, as well as its accumulation in response to both UV-B and HL, suggested a role for LHL4 in photoprotection. To investigate this possibility, we generated *lhl4* knock-out mutants using CRISPR-Cas9 and evaluated their tolerance to HL compared to WT under two distinct physiological conditions: *i.* when grown in LL, without induced photoprotection capacity, and *ii.* when inducing the NPQ capacity through UV-B acclimation, to promote HL tolerance via LHCSR and PSBS (11). Cells from both conditions were subsequently exposed to different light intensities for 1h, and their photosynthetic capacity was tested via maximum quantum yield of PSII (F_v_/F_m_) measurements. Lower F_v_/F_m_ values indicate PSII damage via photoinhibition (46). We confirmed that cells acclimated to UV-B exhibited higher tolerance to HL stress than non-UV-B-acclimated ones (11), as indicated by the higher F_v_/F_m_ values of the former upon 1 h HL stress (compare data in Fig. S5A and S5B). In non-UV-B-acclimated cells, *lhl4* mutants exhibited increasing sensitivity to HL compared to the WT (Fig. 3A) but this difference was almost supressed by UV-B acclimation (Fig. 3B and S5B). We interpret this difference as the consequence of the accumulation of NPQ proteins that could compensate for the absence of LHL4-mediated photoprotection (Fig. 1A). Indeed, the NPQ capacity was similar in WT and *lhl4* cells in UV-B acclimated cells, whereas it remained very low under non-acclimatory LL conditions (Fig. S5C and S5D). Consistently, the Fv/Fm difference between non-acclimated *lhl4* and WT disappeared if the duration of HL exposure increased, i.e. when cells acquire their NPQ capacity through HL exposure and accumulation of LHCSRs (3h, Fig. S5A and S5C). HL-stress generally leads to ROS production in photosynthesis, mainly singlet oxygen (^1^O_2_) at the PSII level (7). We measured ^1^O_2_ levels in WT and *lhl4* during HL exposure and observed a significant overaccumulation of ^1^O_2_ in the *lhl4* strain compared to the WT. This phenomenon occurred in non-acclimated cells (Fig. 3C), but not after UV-B acclimation (Fig. 3D). Thus, our data suggest that LHL4 is required for photoprotection to prevent ROS production and photoinhibition under HL. Our previous findings also suggest that, despite the protective effect of UV-B acclimation, the absence of LHL4 affected cells stress resistance under conditions of extremely intense light irradiance (Fig. 3B, 1’500 µmol photons m^-2^ s^-1^). This implies that LHL4 may play a protective role even in cells that have fully developed their NPQ capacity (Fig. S5D). Accordingly, we observed that LL grown WT and *lhl4* cultures were unable to withstand high-intensity light stress (2’000 µmol photons m^-2^ s^-1^), resulting in complete bleaching within 2 h of exposure (Fig. 3E). On the contrary, both WT and *lhl4* cells maintained their green pigmentation for a longer duration when acclimated to UV-B (Fig. 3F). However, under a prolonged exposure of 8 h, *lhl4* cells failed to endure the stress and bleached, whereas the WT cells remained green. Altogether, these results emphasize the important role of LHL4 in priming UV-B-induced photoprotection in Chlamydomonas along with LHCSRs.

**Fig 3:**
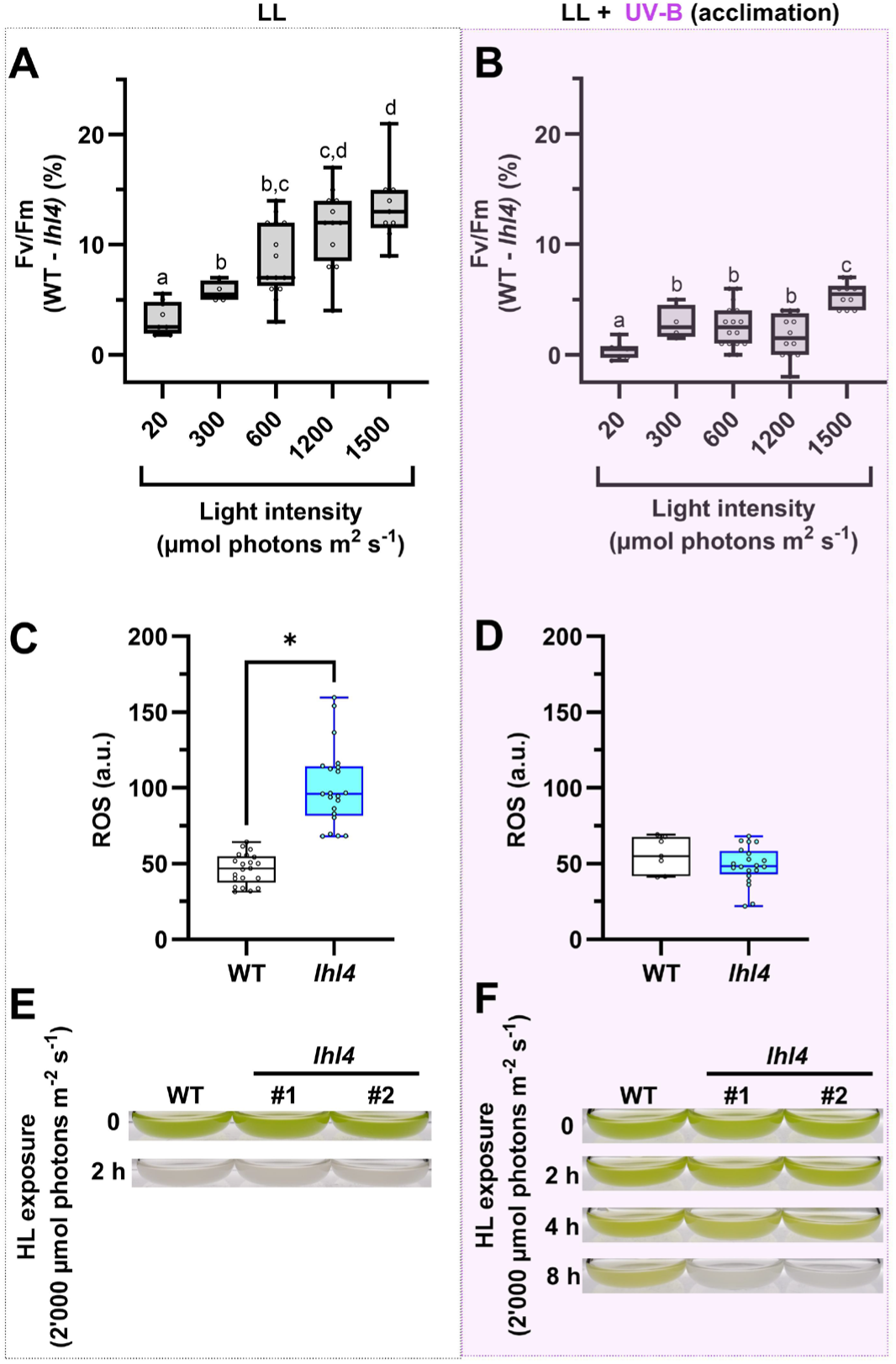
LHL4 mitigates PSII photoinhibition and facilitates cell survival under HL. WT and *lhl4* cells were grown under LL (20 µmol photons m^-2^ s^-1^) **(A**, **C**, **E)** or UV-B acclimated under LL supplemented with UV-B (0.2 mW cm^-2^; LL + UV-B acclimation; **B**, **D**, **F**). **(A-B)** Cells were exposed to different light intensities (20-1500 µmol photons m^-2^ s^-1^) for 1h and placed in the dark for 30 min before assessing their maximum PSII quantum yield (Fv/Fm). For each light intensity, data were calculated as [(WT_Fv/Fm_ - *lhl4*_Fv/Fm_)/(WT_Fv/Fm_)]*100. Different letters above the bars indicate statistically significant difference (p < 0.05); n= 4-16. **(C-D)** WT and *lhl4* cells were exposed to moderate HL (300 µmol photons m^-2^ s^-1^, red light) and ^1^O_2_ accumulation was measured after 30 min of exposure. Asterisk indicates statistical difference (p < 0.01); n= 21. **(E-F)** WT and *lhl4* flasks were photographed after the LL or UV-B acclimation period (0). They were then placed under strong HL (2’000 µmol photons m^-2^ s^-1^) and photographed after 2, 4 and 8 h.

### LHL4 binds the PSII monomer upon UV-B exposure

To elucidate the mechanistic basis underlying LHL4-mediated photoprotection in Chlamydomonas, we performed biochemical analysis under native conditions by Blue Native Polyacrylamide Gel Electrophoresis (BN-PAGE) of solubilized thylakoid complexes isolated from WT cells exposed to UV-B, coupled with crosslinking mass spectrometry (XL-MS) analysis. The latter approach allows capturing protein-protein interaction by creating covalent bonds between amino acids in close proximity and can be directly applied to complexes previously separated via BN-PAGE (47). Importantly, the chemical crosslinking reaction did not produce any noticeable difference on the presence of major photosynthetic complexes (Fig. S6A). We found that LHL4 was only detected upon UV-B exposure and migrated in proximity to the PSII monomer/cytochrome b_6_f bands, as well as in the LHCII trimer region (Fig. 4A). XL-MS data acquired on these specific gel bands were first validated by mapping the crosslinks on known PSII and cytochrome b_6_f structures (Fig. S6B-D), where most fall within an acceptable 20 Å distance cut-off (Fig. S6C) (48). Although LHL4 seemed sub-stoichiometric when compared to PSII and cytochrome b_6_f subunits, we found it uniquely associated with the PSII core antenna. This conclusion is corroborated by simulations of the pairwise interactions LHL4-CP43 (Fig. S7) and LHL4-CP47 (Fig. S8) using AlphaFold2 multimer (47). We therefore propose that LHL4 binding to the PSII monomer during exposure to HL promotes the formation of an intermediate shielded form that protects the complex from photoinhibition. The structural mechanism is reminiscent of that recently proposed for the PSII oxygen-evolving complex, where the Ycf48 assembly factor binds to the PSII monomer, promoting the initial steps of PSII assembly in cyanobacteria (49).

**Fig 4:**
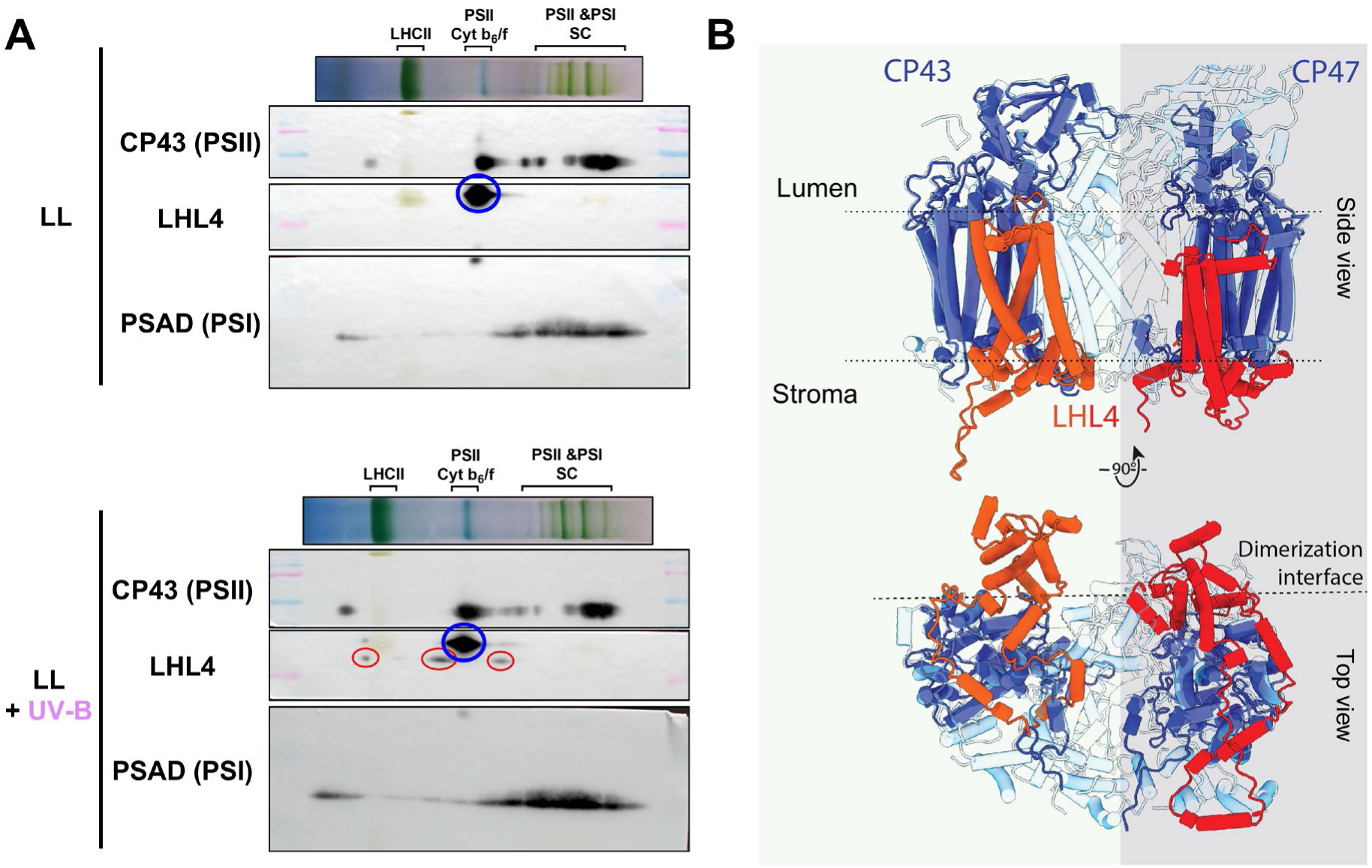
LHL4 localizes with PSII monomers at the dimerization interface interacting with the inner PSII antenna proteins CP43 and CP47. **(A)** BN-PAGE of thylakoids extracted from cells exposed to LL (20 µmol photons m^-2^ s^-1^) or LL supplemented with UV-B (LL +UV-B; 0.2 mW cm^-2^). Second dimension blots indicate LHL4 (red circles), and its absence in LL cells. A blue circle indicates a non-specific band detected by the LHL4 antibody in both WT and *lhl4*. **(B)** LHL4 interactors CP43 and CP47 uncovered by crosslinking and in-gel digestion of the BN-PAGE band. The two AlphaFold2 pairwise predictions of LHL4-CP43/47 are shown overlaid on the PSII monomer structure (PDB 6KAD, light blue) and both are localized at the dimerization interface.

## Conclusion

Our study reveals a key role for the LHL4 protein in photoprotection. LHL4 protects PSII during both the early steps of photodamage and steady state conditions by associating with the core complex of the PSII monomer via two antenna proteins, CP43 and CP47. Photoprotection occurs upon exposure to UV-B and HL, which concomitantly activate the UVR8 and PHOT photoreceptor responses thanks to an early convergence of the two photoreceptors signaling pathways likely at the COP1 level. Our phylogenetic analysis indicates that LHL4 is exclusively present in green microalgae. However, its crucial function in photoprotection of these organisms suggests the existence of similar actors in other phototrophs. Consistently, specific High-Light-Inducible Proteins (HLIPs) have been identified in cyanobacteria, where they associate with CP47 and intermediates of PSII assembly modules (50). Remarkably, cyanobacteria lacking HLIPs are viable but experience extreme light-induced stress levels, suggesting the presence of protective mechanism in prokaryotic photosynthetic organisms similar to the one conferred by Chlamydomonas LHL4. Investigating potential candidates in other microalgae and flowering plants may unlock a deeper understanding of photoprotection mechanisms across diverse photosynthetic life forms.

## Materials and Methods

### Algae Strains

Chlamydomonas mutant strains *lhl4* (LHL4^K34*^; * indicating a translation stop codon), *uvr8* (UVR8^S86*^) and *phot* (PHOT^C32*^) were generated in the WT137C parental strain (51) (Fig. S9) by introducing a TAA STOP codon by homology-directed repair using CRISPR-Cas9 (52, 53). The following target sequences were used: 5’-CGTGTCAACGCCGAGAAAAG-3’ for *lhl4*, 5’-CGAGGACAGATCTACAGCTG-3’ for *uvr8* and 5’-CACGCTTCCGGACTGTCCGC-3’ for *phot.* The following templates for DNA repair were used: 5’-CGCAGTGATTGCCCGTGTCAACGCCGAGAATATAAGGACCACGACATCGACTACAAGGACAAGTGGCTTTGCAAAGGTAAGCTGAGAACT-3’ for *lhl4*, 5’-GGTCGCGTCGTCACGAGGACAGATCTACAGTATAAGGACCACGACATCGACTACAAGGACCTGGGGCTGGTGGGTGGCGTTGGGGCGCAG-3’ for *uvr8* and 5’-TCGTCGCAGATGCCACGCTTCCGGACTGTCTATAAGGACCACGACATCGACTACAAGGACCGCTGGTCTACGCCAGCGAGGGGTGAGCGG-3’ for *phot*.

The *cop1^hit1^* (32) was used for comparison with the WT CC124. *Crco* (LMJ.RY402.149321) (27, 31) and *crblz3* (LMJ.RY402.194448) (38) mutants were obtained from the Chlamydomonas Library Project (CLiP) (54) together with their corresponding WT (CC5325). *uvr8* (LMJ.RY40202.156289) from the CLiP library was used for the RNA-seq experiment (11).

### Algae culture and light treatment

Algae cells were grown in the Tris Acetate Phosphate (TAP) growth medium (55) under continuous light (20 µmol photons m^-2^ s^-1^, from LED panels), 25°C, and shaking (110 rpm). In all experiments, cells were harvested during the exponential phase (2-3 x 10^6^ cells per mL), washed, and resuspended in High Salt Medium (HSM) minimal growth medium at 4.5 x 10^6^ cells per mL. They were placed for at least 1 h under dim light (20 µmol photons m^-2^ s^-1^) before applying specific light treatments. UV-B-exposed samples (+ UV-B) were treated by Philips TL20W/01S narrowband UV-B tubes filtered with a WG filter (Schott Glaswerke) with half-maximal transmission at 311 nm (0.07 or 0.2 mW cm^2^) (56). Control (-UV-B) samples were placed under a WG filter with half-maximal transmission at 360 nm to block UV-B. In both conditions, samples were concomitantly exposed to dim light (20 µmol photons m^-2^ s^-1^) provided by fluorescent white light tubes (Osram Dulux L). Cells were exposed to UV-B for 6 h, with the exception of Figure 1A, 3G and 3H, where the treatment duration was extended to 16 h to fully induce the UV-B protection of the photosynthetic machinery, as detailed in (11). For HL treatment, cells were placed under LED panels (SL 3500, Photon Systems Instruments) at the indicated light intensity.

### Phylogenetic analysis

The sequences of LHL4, PSBS1, LHCSR3.1, and ELIP1-9 were subjected to a blastp analysis using the online tool available at https://www.ncbi.nlm.nih.gov/ with a percent identity range of 35-100% and a query coverage range of 50-100%. Protein sequences were then aligned using the MAFFT tool (auto strategy, https://mafft.cbrc.jp/alignment/server/). To enhance the accuracy of the alignment, manual curation was performed using Jalview to eliminate any duplicate or misaligned sequences. The phylogenetic tree was constructed using IQ-TREE, employing the LG+F+G4 model. The generated tree was then visualized graphically using iTOL v6, a web-based tool accessible at https://itol.embl.de/.

### Protein extraction and immunoblot analysis

Cells were pelleted and resuspended in 80% (v/v) acetone. Proteins were precipitated by centrifugation at 4°C and resuspended in lysis buffer (100 mM Tris-HCl, pH 6.8, 4% (v/v) SDS, 20 mM EDTA) supplemented with protease inhibitor (cOmplete, Roche). 10-30 µg of proteins were separated by SDS-PAGE in Tris-Glycine buffer (25 mM Tris, 190 mM glycine, 0.05 % SDS), transferred onto nitrocellulose membrane using the same buffer supplemented with 20 % (v/v) ethanol for 80 minutes at 110 V. Anti-CrUVR8 (24), anti-PHOT (16), anti-LHCSR3 (AS142766; Agrisera), anti-LHCSR1 (AS142819; Agrisera), anti-LHL4 (AS07250, Agrisera), anti-ATPB (AS05085; Agrisera), anti-CP43 (AS111787; Agrisera), anti-PSAH (AS06143; Agrisera) were used in this study.

### RNA extraction

20 million cells were pelleted and frozen in liquid nitrogen. RNA were extracted using the RNeasy Plant Mini Kit (Qiagen) including a DNase treatment to remove residual genomic DNA (RNase-Free DNase Set, Qiagen).

### Quantitative Real-Time PCR (RT-qPCR)

cDNA synthesis was performed with the TaqMan Reverse Transcription Reagents kit (Applied Biosystems). Amplification by RT-qPCR was performed using cDNA in the presence of SYBR Green (Master Mix PCR Power SYBR^TM^, Applied Biosystems) and specific primers (0.3 µM) for the amplification of the gene of interest, using a CFX Connect Real-Time PCR Detection System (Biorad). Data was analyzed using the ΔΔCt method (57) and the Cre06.g278222 reference gene (24, 58). The following primers were used: 5’-TACGGTGTGGATGACGTGAC-3’ and 5’-AGGTGATAATCTGGCGGATG-3’ for *LHL4*, and 5’-CTTCTCGCCCATGACCAC-3’ and 5’-CCCACCAGGTTGTTCTTCAG-3’ for Cre06.g278222.

### RNA-seq analysis

Cells from WT CC-5325, *uvr8*, *crco* and *crblz3* were prepared as described above (“RNA extraction”) and then exposed for 1 h to LL (20 µmol photons m^-2^ s^-1^; -UV-B control samples) or LL supplemented with 0.2 mW cm^-2^ of UV-B (+UV-B). Total RNA was extracted from three independent biological replicates. The RNA quality control, library preparation using TruSeqHT Stranded mRNA (Illumina) and sequencing on an Illumina HiSeq 4000 System using 100-bp single-end reads protocol were performed at the iGE3 genomics platform of the University of Geneva. Quality control was performed with FastQCv.0.11.9. Reads were mapped to the Chlamydomonas genome CreinhardtiiCC_4532_707_v6.1 (PhytozomeV13) using STARv.2.7.10b software (59), with average alignment of 91.65%. Raw counts were obtained using HTSeq v.0.11.3 (60). Filtering out lowly expressed genes (13’613 genes were kept), normalization and differential expression analysis were performed with the R/Bioconductor package edgeR v.3.42.4 (61), and statistical significance was assessed with a general linear model, negative binomial distribution, and quasi-likelihood F test. Genes with a fold change ≥ 2 and p-value ≤ 0.05 (with a multiple testing Benjamini and Hochberg FDR correction) were considered differentially expressed. Annotations (v6.1) were obtained from Phytozome database (v13). The RNA-Seq data reported have been deposited in the NCBI Gene Expression Omnibus (www.ncbi.nlm.nih.gov/geo).

### ROS quantification

^1^O_2_ production was estimated using the Singlet Oxygen Sensor Green (SOSG, Invitrogen) dye (62). Cultures at 4.10^6^ cells/mL were placed in 24-wells plates. 0.5 µM of SOSG were added to each well, before exposure for 30 min to red light (630 nm, 300 µmol photons m^-2^ s^-1^). This wavelength should minimize photobleaching of the dye, which is sensitive to blue light and prolonged exposure to high-intensity illumination (63). The fluorescence of each well was then measured with a plate reader (Infinite M1000; Tecan) (λ excitation: 504 nm; λ emission: 525 nm ± 10 nm).

### NPQ and photoinhibition measurements

Chlorophyll fluorescence was imaged in 100 µl of cell suspension placed on a 96-well plate. Cells were dark-adapted for 10 min before measurement. Fluorescence was quantified using a Speedzen III fluorescence imaging setup (JBeam Bio). Maximum fluorescence in the dark (F_m_) or during actinic light exposure (F_m’_, 900 µmol photons m^-2^ s^-1^) was measured using saturating red pulses (250 ms, 3’000 µmol photons m^-2^ s^-1^) followed by blue light (470 nm) detection pulses (10 µs). NPQ was calculated as (F_m_-F_m’_)/F_m’_ (64). For photoinhibition experiments, cells were treated for the indicated times with HL and then dark adapted for 30 min. The chlorophyll fluorescence was then measured before (F_0_) and during the saturating pulse (F_m_), and the maximum quantum yield of PSII was calculated as F_v_/F_m_=(F_m_-F_0_)/F_m_.

### Mass spectrometry (MS)-based proteomic analyses

Three biological replicates of Chlamydomonas cells exposed or not to UV-B for 16 h were prepared. The Chlamydomonas cells were disrupted in a tube containing glass beads using a Precellys instrument (Bertin Technologies) at 7’500 rpm and 4°C for two cycles of 30-sec each, with a 30-sec pause in between. Subsequently, the samples were centrifuged at 20’000g for 5 min at 4°C. The resulting pellets, enriched with membrane proteins, were resuspended in a solution containing 50 mM Tris-HCl pH 6.8, 2% SDS, 10 mM EDTA, and protease inhibitor (cOmplete, Roche) to maintain the integrity of the proteins during further processing. Proteins were then solubilized in buffer (65) and heated for 10 min at 95°C. They were then stacked in the top of a 4-12% NuPAGE gel (Invitrogen), stained with Coomassie blue R-250 (Bio-Rad) before in-gel digestion using modified trypsin (Promega, sequencing grade) as previously described (66). The resulting peptides were analyzed by online nanoliquid chromatography coupled to MS/MS (Ultimate 3000 RSLCnano and Q-Exactive HF, Thermo Fisher Scientific) using a 120-min gradient. For this purpose, the peptides were sampled on a precolumn (300 μm x 5 mm PepMap C18, Thermo Scientific) and separated in a 75 μm x 250 mm C18 column (Reprosil-Pur 120 C18-AQ, 1.9 μm; Dr. Maisch GmbH). The MS and MS/MS data were acquired using Xcalibur version 2.9 (Thermo Fisher Scientific).

Peptides and proteins were identified by Mascot (version 2.8.0, Matrix Science) through concomitant searches against the Chlre5_6 database (downloaded from JGI Genome Portal, (67), 19’526 sequences), the mitochondrion and chloroplast protein sequences (downloaded from NCBI, respectively 69 and 8 proteins), and a homemade database containing the sequences of classical contaminant proteins found in proteomic analyses (human keratins, trypsin, … 126 sequences). Trypsin/P was chosen as the enzyme and two missed cleavages were allowed. Precursor and fragment mass error tolerances were set at respectively at 10 and 20 ppm. Peptide modifications allowed during the search were: Carbamidomethyl (C, fixed), Acetyl (Protein N-term, variable) and Oxidation (M, variable). The Proline software (version 2.2.0, (68)) was used for the compilation, grouping, and filtering of the results (conservation of rank 1 peptides, peptide length ≥ 6 amino acids, FDR of peptide-spectrum-match identifications < 1% (69), and minimum of one specific peptide per identified protein group). Proline was then used to perform a MS1 label-free quantification of the identified protein groups based on razor and specific peptides.

Statistical analysis was performed using the ProStaR software (70) based on the quantitative data obtained with the three biological replicates analyzed per condition. Proteins identified in the contaminant database, proteins identified by MS/MS in less than two replicates of one condition, and proteins quantified in less than three replicates of one condition were discarded. After log2 transformation, abundance values were normalized using the variance stabilizing normalization (vsn) method, before missing value imputation (SLSA algorithm for partially observed values in the condition and DetQuantile algorithm for totally absent values in the condition). Statistical testing was conducted with limma, whereby differentially expressed proteins were selected using a log_2_FC cut-off of 1 and a p-value cut-off of 0.00776, allowing to reach a FDR inferior to 1% according to the Benjamini-Hochberg estimator. Proteins found differentially abundant but identified by MS/MS in less than two replicates or detected in less than three replicates in the condition in which they were found to be more abundant were manually invalidated (p-value = 1).

### Thylakoids isolation and BN-PAGE

400 million cells were broken in a tube containing glass beads using a Precellys instrument (Bertin Technologies) at 7’500 rpm and 4°C for two cycles of 30-sec each, with a 30-sec pause in between. Subsequently, the samples were centrifuged at 20’000g for 5 min at 4°C. After centrifugation (20’000 g, 4°C, 5 min), the pellet was resuspended in 25 mM Hepes pH 7.5, 5 mM MgCl_2_, 0.3 M sucrose, and protease inhibitor (cOmplete, Roche). Thylakoids were isolated using a discontinuous three-step sucrose gradient (2.3 M, 1.3 M, 0.5 M) after centrifugation (76’000 g, 1 h, 4°C). The chlorophyll concentration was estimated, and 20 µg of chlorophyll was solubilized in 1% α-dodecylmaltoside for 5 min at room temperature. For the BN-PAGE analysis, a loading buffer (0.5 M aminocaproic acid, 30% sucrose, 100 mM bistris HCl pH 7, and 50 mg/mL blue Coomassie G-250) was added to the sample. The thylakoid extract was then loaded onto a 4-16% NativePAGE (InVitrogen), and gel electrophoresis was conducted with increasing voltage intensity (ranging from 75 V to 200 V) in an anode buffer (50 mM Bis-Tris HCl pH 7) and a blue cathode buffer (50 mM Tricine, 15 mM Bis-Tris HCl pH 7, 0.01% blue Coomassie G-250). When the migration reached the middle of the gel, the blue buffer was replaced with fresh cathode buffer devoid of Coomassie blue.

### Crosslinking MS and structural modelling

UV-B treated and un-treated (control) cells were disrupted in a tube containing glass beads using a Precellys instrument (Bertin Technologies) at 7’500 rpm and 4°C for two cycles of 30-sec each, with a 30-sec pause in between. Cell lysates were chemically crosslinked with 8 mM 4-(4,6-dimethoxy-1,3,5-triazin-2-yl)-4-methyl-morpholinium chloride (DMTMM) for 30 min at 4°C in the darkness). We used the water soluble DMTMM reagent because of the scarce availability of exposed Lys residues in the loops of LHL4. DMTMM is considered as a “short-range” crosslinker, lacking a spacer arm and with reactive groups targeting carboxylic acids and primary amines (aspartic and glutamic acids to lysine and amino termini of proteins). This reagent provides distance constraints between 0 and ∼25 Å considering the flexibility of the side-chains (7 and 5 Å for lysine and carboxylic acids, respectively), the α-carbon backbone (6 Å) and the overall protein flexibility (48, 71). Crosslinked thylakoids were purified according to established protocols and then solubilised for BN-PAGE as described above. Comparison of crosslinked and non –crosslinked thylakoids showed no noticeable differences in the band patterns, we thus assumed the XL reaction did not produce artefacts (Fig. S5). Gel bands containing LHL4 from 3 gel lanes (20 µg of Chl per lane) were excised and pooled. Additionally, a second XL reaction with 16 mM DMTMM for 30 min at room temperature was done on the excised bands and reaction was quenched by addition of 1 M Tris HCl to a final concentration of 10 mM. Gel bands were then in-gel digested (66) and approximately 200 ng injected onto a LC-MS/MS set-up similar as for the proteomic analysis, with minor modifications. Briefly, LC separation gradients were of 180 min with elution gradient profiles as follows: 0 – 10% solvent B (0.1% (v/v) formic acid in 80% (v/v) acetonitrile) over 10 min, 12 - 35% solvent B over 125 min, 36 - 44% solvent B over 20 min, 45-100% solvent B over 10 min, and finally 100% B for 15 min. Full-scan MS spectra were collected in a mass range of m/z 350 – 1300 Th in the Orbitrap at a resolution of 60’000 at m/z=200 Th after accumulation to an AGC (Automatic Gain Control) target value of 1e6 with a maximum injection time of 50 ms. In-source fragmentation was activated and set to 15 eV. The cycle time for the acquisition of MS/MS fragmentation scans was set to 2 s. Charge states accepted for MS/MS fragmentation were set to 3 - 8. Dynamic exclusion properties were set to n = 1 and to an exclusion duration of 15 s. Stepped HCD (Higher energy Collision Dissociation) fragmentation (MS/MS) was performed with increasing normalized collision energy (27, 30, 33 %) and the mass spectra acquired in the Orbitrap at a resolution of 30,000 at m/z=200 Th after accumulation to an AGC target value of 1e5 with an isolation window of m/z = 1.4 Th and maximum injection time of 120 ms. Additionally, LHL4-containing BN-PAGE bands were acquired with a MS method similar to that used for normal tryptic peptides as described above in order to screen proteins present in the BN-PAGE band.

Raw data were searched using pLink2 (72) for crosslinked peptide pairs or as described above for normal peptide search. A minimal peptide length of six and two miss cleaved sites was allowed. Cysteine carbamidomethylation was set as fixed modification. Methionine oxidation, protein N-term acetylation and lysine acetylation were set as dynamic modifications. For crosslinked peptides, the reference database containing only the proteins identified from the normal peptide was used, but with an increased number of missed cleavages allowed of 3. Identified crosslinks were only accepted through at 1% FDR filter and if present in both replicates.

Crosslinks identified were mapped using the ChimeraX plugin XMAS (73) on the PSII monomer (extrapolated from PDB #6KAD) and Cyt b6f (PDB #1Q90) structures. The predicted LHL4 AlphaFold2 model downloadable from UniprotKB was used. For additional validation of the interaction we docked in parallel LHL4 for the core complex on the HADDOCK webserver (74), using the identified crosslinks as distance constraints and predicting the pairwise interaction between CP43/CP47 and LHL4 with Alphafold2 multimer algorithm in ColabFold (75).

## Acknowledgements

We thank Olaf Kruse for kindly providing the *cop1^hit1^* mutant and Michel Goldschmidt-Clermont and Stéphane Ravanel for helpful discussions throughout the project. We thank Anja Krieger-Liszkay for technical discussions regarding ROS measurement, and Caroline Juery, Yamama Naciri and Charles Pouchon for their assistance in conducting the phylogenetic analysis of LHL4. We also thank Florence Courtois, Gilles Curien and Michel Goldschmidt-Clermont for their critical reading of the manuscript. The RNA-Seq experiments were performed at the iGE3 Genomics Platform of the University of Geneva (https://ige3.genomics.unige.ch/), and we thank Natacha Civic and Céline Delucinge Vivier for bioinformatic analysis of the RNA-Seq data. This work was supported by an IDEX Université Grenoble Alpes International Strategic Partnership grant (SUnsPOT), the University of Geneva, and, in part, by the Swiss National Science Foundation (grant 310030_207716 to R.U.). P.A. acknowledges funding from the European Union’s Horizon 2020 research and innovation programme under the Marie Skłodowska-Curie grant agreement No 101066400 - PHOTO-LINK. C.G. and G.F. acknowledge funding from the European Research Council ERC (Chloro-Mito; grant no. 833184). G.A. and C.B. acknowledge funding from the CNRS Momentum program. This work is also supported by the French National Research Agency in the framework of the “Investissements d’avenir” program (ANR-15-IDEX-02) and by GRAL, a program from the Chemistry Biology Health (CBH) Graduate School of University Grenoble Alpes (ANR-17-EURE-0003) (E.C., Y.C., G.F. and G.A.). Proteomic experiments were partially supported by Agence Nationale de la Recherche under projects ProFI (Proteomics French Infrastructure, ANR-10-INBS-08).

## Author contributions

M.D., G.F., E.D., R.U. and G.A. designed research; M.D., C.B., E.C., P.A., D.T., C.G., M.M., S.B., E.D. and G.A. performed research; M.D., P.A., D.T., S.B., Y.C., G.F., E.D., R.U. and G.A. analyzed data; and G.F., E.D., R.U. and G.A. wrote the paper with inputs of all coauthors.

## Data availability

All MS raw data files, including the search results (spectral matches and the crosslink tables), are deposited to the ProteomeXchange Consortium via the PRIDE partner repository. The RNA-Seq data reported in this article have been deposited in the NCBI Gene Expression Omnibus.

## Accession numbers

Sequence data from this article can be found in the Joint Genome Institute Phytozome data libraries (https://phytozome.jgi.doe.gov/pz/portal.html) under the accession numbers *LHL4*, Cre17.g740950. *UVR8*, Cre05.g230600. *PHOT*, Cre03.g199000. *COP1*, Cre02.g085050. *CrCO*, Cre06.g278159. *CrBLZ3*, Cre06.g310500. *LHCSR1*, Cre08.g365900. *LHCSR3*, Cre08.g367500 and Cre08.g367400. *PSBS*, Cre01.g016600 and Cre01.g016750, *ELIP1*, Cre14.g626750. *ELIP2*, Cre16.g679250. *ELIP3*, Cre02.g143550. *ELIP4*, Cre07.g320400. *ELIP5*, Cre04.g211850. *ELIP6*, Cre13.g576760. *ELIP7*, Cre08.g384650. *ELIP8*, Cre09.g393173. *ELIP9*, Cre09.g394325. *RACK1*, Cre06.g278222.

---

**Fig S1:**
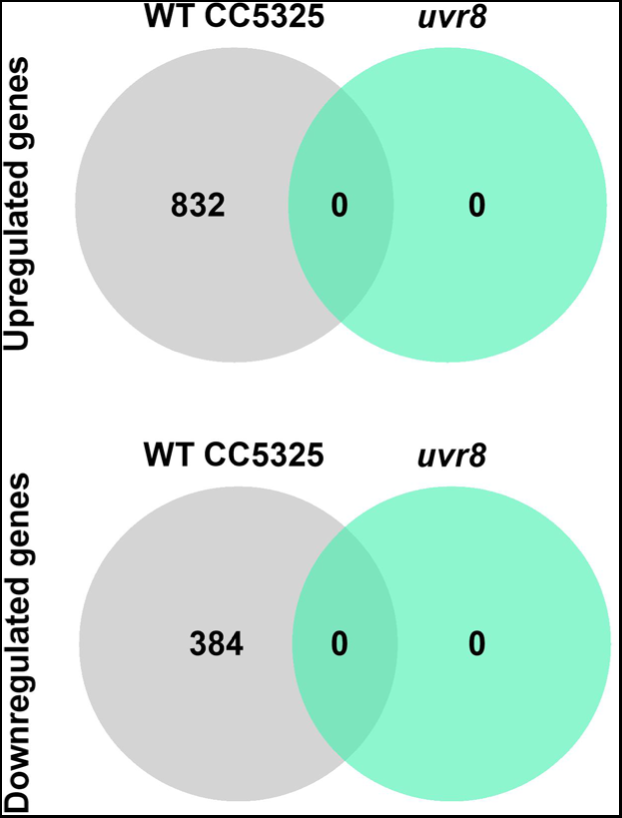
Transcriptome analysis of WT and *uvr8* in response to UV-B. Venn diagrams showing the number of genes differentially regulated in LL + UV-B versus LL (downregulated genes: log_2_FC≤-1, upregulated genes: log_2_FC≥1; FDR≤0.05; n=3) in WT strain CC5325 and *uvr8* null mutant.

**Fig S2:**
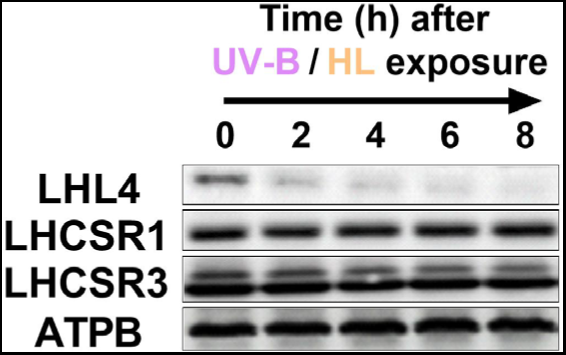
Comparative stability of LHL4 and LHCSRs proteins. Immunodetection of LHL4, LHCSR1, LHSCR3 and ATPB in cells sequentially exposed to LL (20 µmol photons m^-2^ s^-1^) supplemented with UV-B (0.2 mW cm^-2^) for 16 h and then HL (300 µmol photons m^-^² s^-1^) for 4 h. Cells were finally placed under LL (20 µmol photons m^-2^ s^-1^) for the indicated times. ATPB was used as loading control.

**Fig S3:**
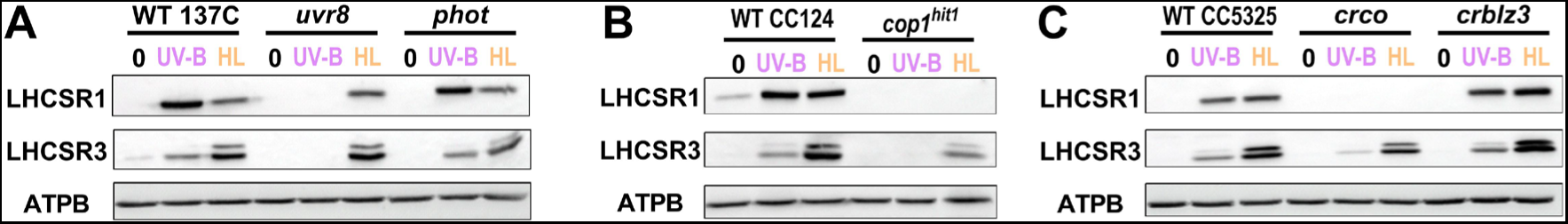
Accumulation of LHCSR1 and LHCSR3 in mutants with impaired UV-B and/or HL signaling pathways. **(A-C)** Immunodetection of LHCSR1 and LHCSR3 in cells grown under LL (0), exposed for 6 h to LL supplemented with UV-B (0.2 mW cm*^-^*^2^, UV-B) or HL (300 µmol photons m^-2^ s^-1^, HL). Accumulation was assessed in **(A)** *uvr8* and *phot*, **(B)**, *cop1^hit1^* and **(C**) *crco* and *crblz3* **(C)** ATPB was used as loading control.

**Fig S4:**
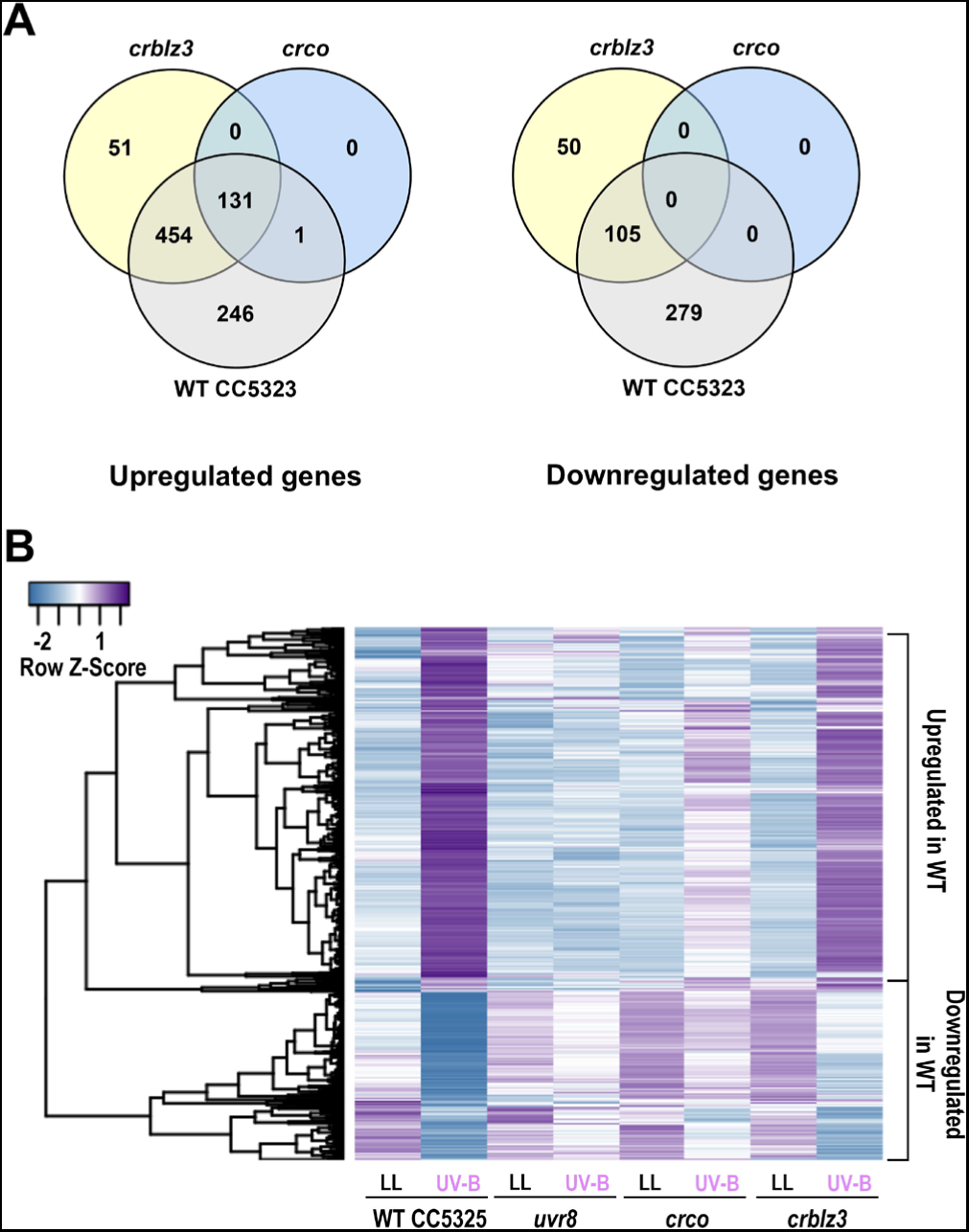
Transcriptome analysis of WT and *uvr8*, *crco* and *crblz3* mutants in response to 1h UV-B. **A)** Venn diagrams indicating the number of genes differentially regulated in LL + UV-B versus LL (left panel: induced genes (UP) with log_2_FC≥1 and FDR≤0.05; right panel: repressed genes (DOWN) with log_2_FC≤1 and FDR≤0.05; n=3); and the intersection between the genotypes. **B)** Hierarchical clustering of normalized expression patterns (Z-score) in WT, *uvr8*, *crco* and *crblz3* of genes differentially regulated in WT LL + UV-B versus LL (log2FC≤1 or ≥1, FDR≤0.05).

**Fig S5:**
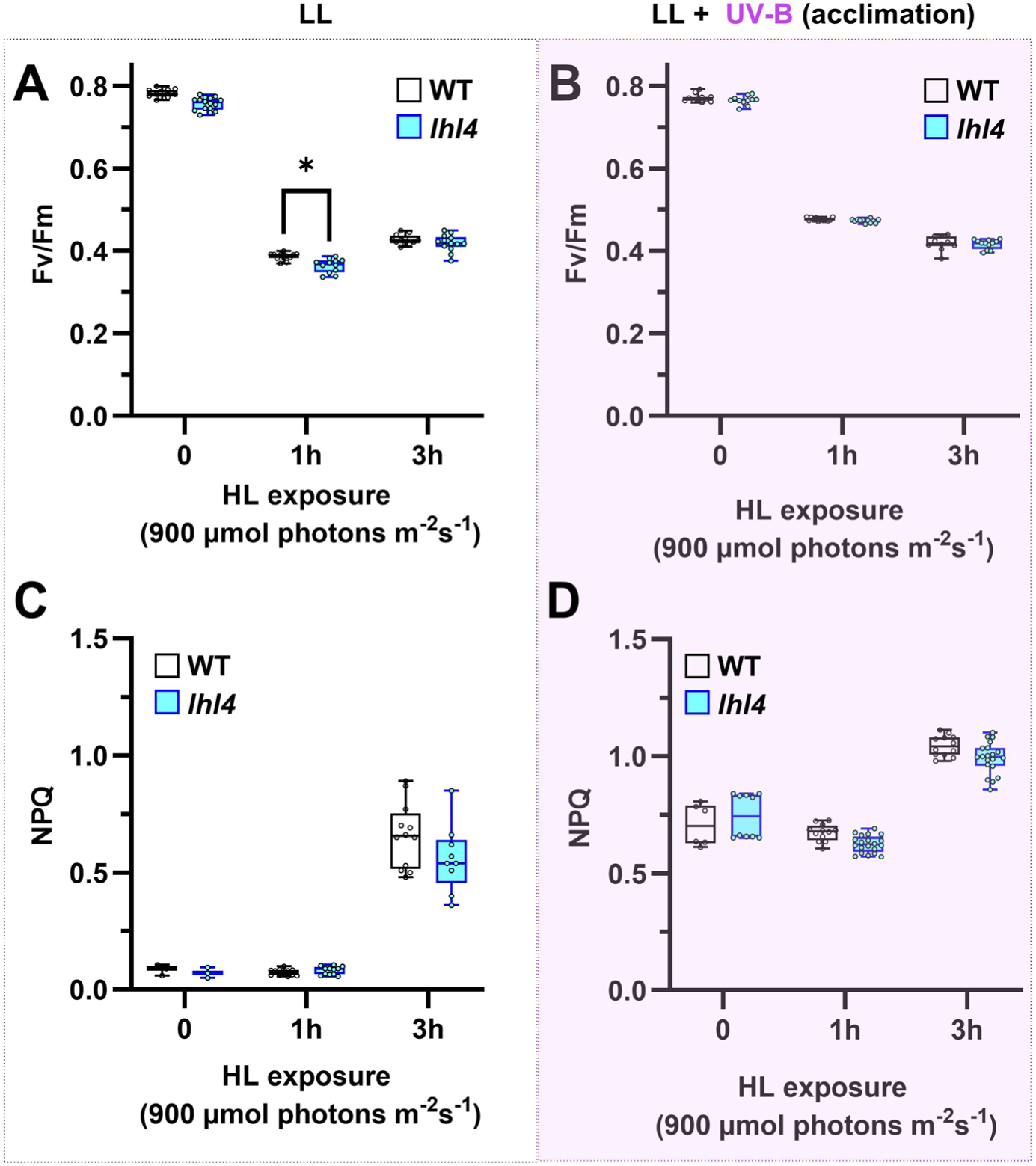
UV-B acclimation protects *lhl4* from photoinhibition. WT and *lhl4* cells were grown under LL (20 µmol photons m^-2^ s^-1^) **(A**, **C)** or under LL supplemented with UV-B (0.2 mW cm^-2^; LL + UV-B acclimation) **(B**, **D**). Cells were exposed to HL (900 µmol photons m^-2^ s^-1^) for 1h or 3h and placed in the dark for 30 min. Fv/Fm and NPQ were then evaluated using chlorophyll fluorescence measurements. Asterisk indicates statistical significant difference between WT and *lhl4* (p < 0.01); n= 7-12.

**Fig S6:**
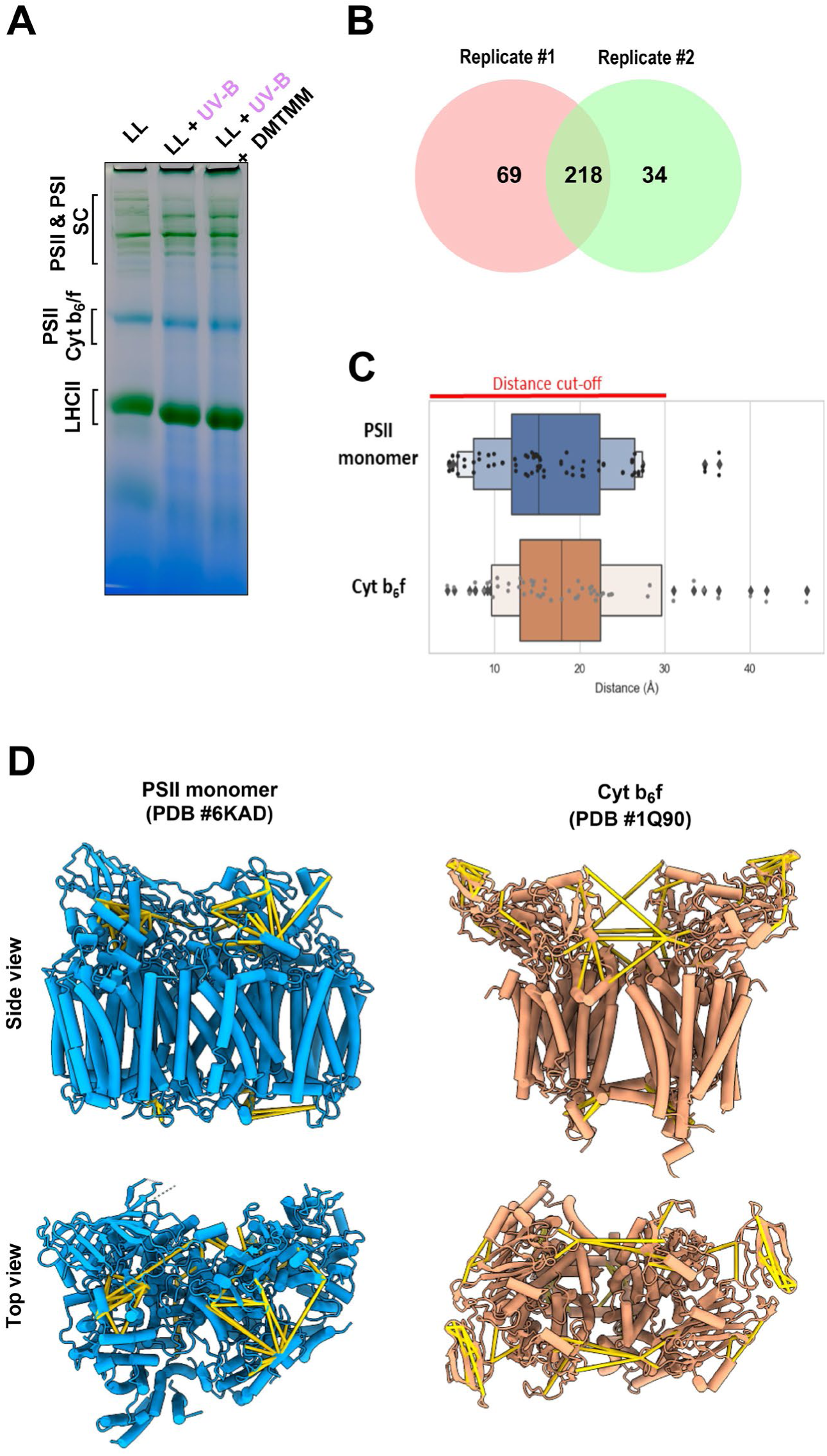
In-gel crosslinking MS does not produce structural artefacts. **(A)** BN-PAGE of thylakoids extracted from WT cells grown under LL (20 µmol photons m^-2^ s^-1^) or UV-B acclimated under LL supplemented with UV-B (0.2 mW cm^-2^; LL + UV-B) in the presence or absence of the crosslinker (DSSO). **(B)** Upon in-gel digestion 218 reproducible crosslinks from two independent replicates could be identified. **(C)** As a quality check, plotted crosslinks fall within the acceptable distance range of 30 Å between Cα-Cα of the crosslinked lysine residues, with the vast majority within the strict cut-off of 20 Å. **(D)** The identified crosslinks (yellow) were plotted onto known structures of Chlamydomonas PSII monomer (PDB #6KAD, left) and cytochrome b_6_f (PDB #1Q90, right).

**Fig S7:**
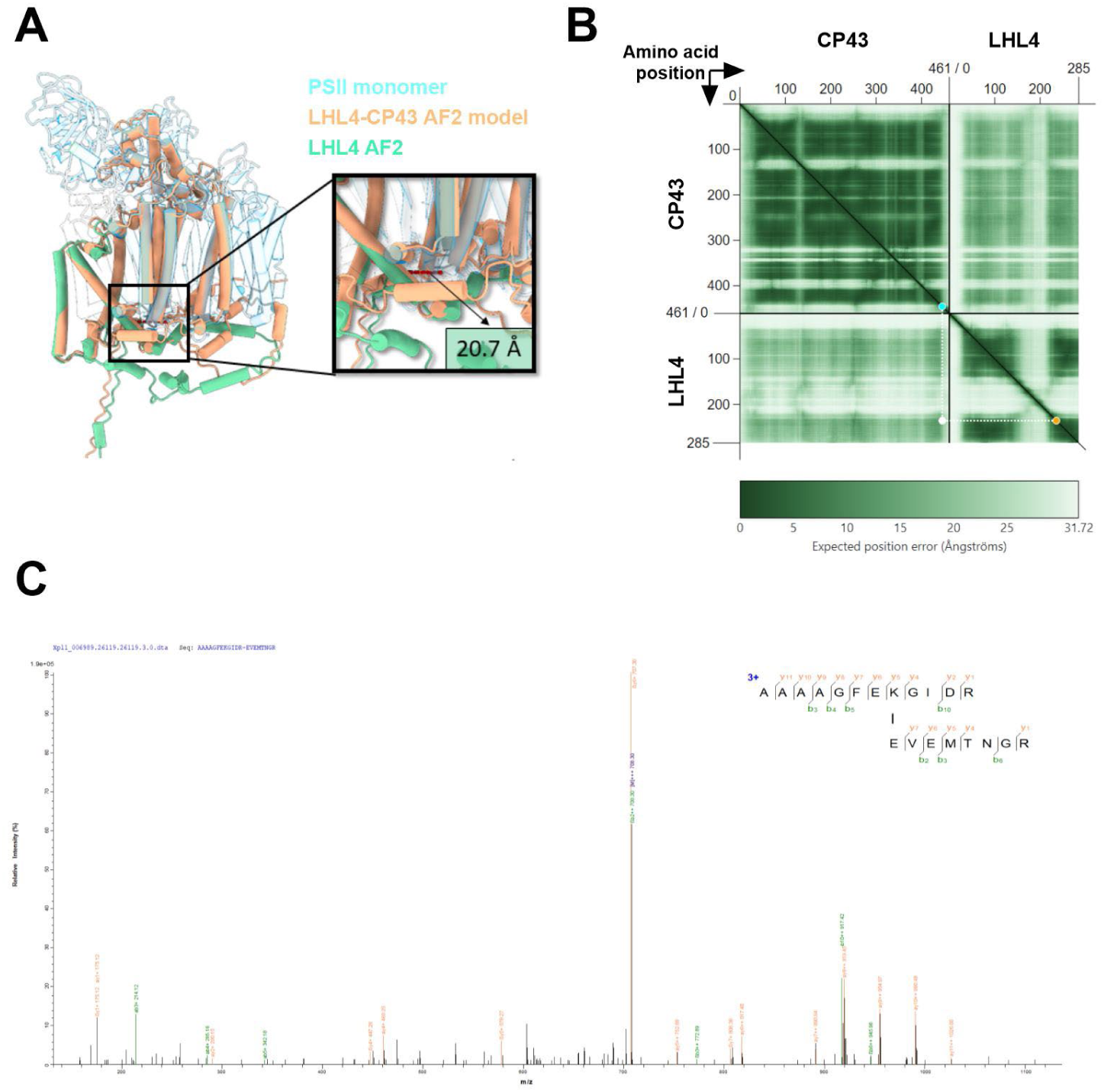
LHL4-CP43 predicted complex supported by experimentally validated crosslinks. **(A)** LHL4-CP43 interaction modelled with AlphaFold2 (AF2) multimer and the crosslinks distance mapped on the model**. (B)** Predicted Aligned Error (PAE) plot indicating the expected positional error between and within each residue of CP43 and LHL4. The detected crosslink is represented as the intersection (white dot) between LHL4-Glu238 (orange dot) and CP43-Lys445 (blue dot). **(C)** Representative fragmentation spectra of the experimentally derived crosslink between LHL4 and CP43.

**Fig S8:**
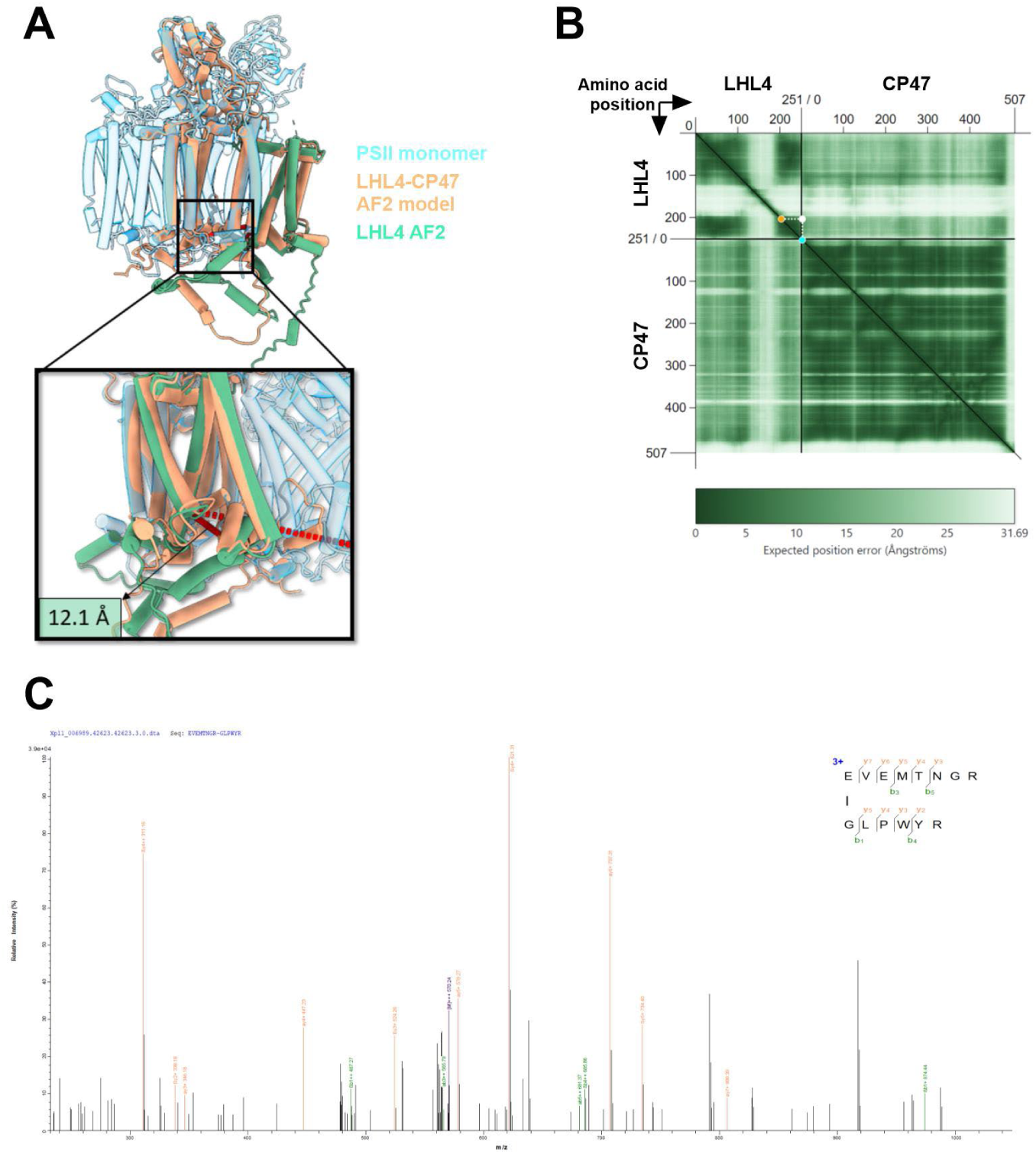
LHL4-CP47 predicted complex supported by experimentally validated crosslinks. **(A)** LHL4-CP43 interaction modelled with AlphaFold2 (AF2) multimer and the crosslink distance showed on the model. **(B)** Predicted Aligned Error (PAE) plot indicating the expected positional error between and within each residue of protein CP47 and LHL4. The detected crosslink is represented as the intersection (white dot) between LHL4-Glu238 (orange dot) and CP47 N-terminus (blue dot). **(C)** Representative fragmentation spectra of the experimentally derived crosslink between LHL4 and CP47.

**Fig S9:**
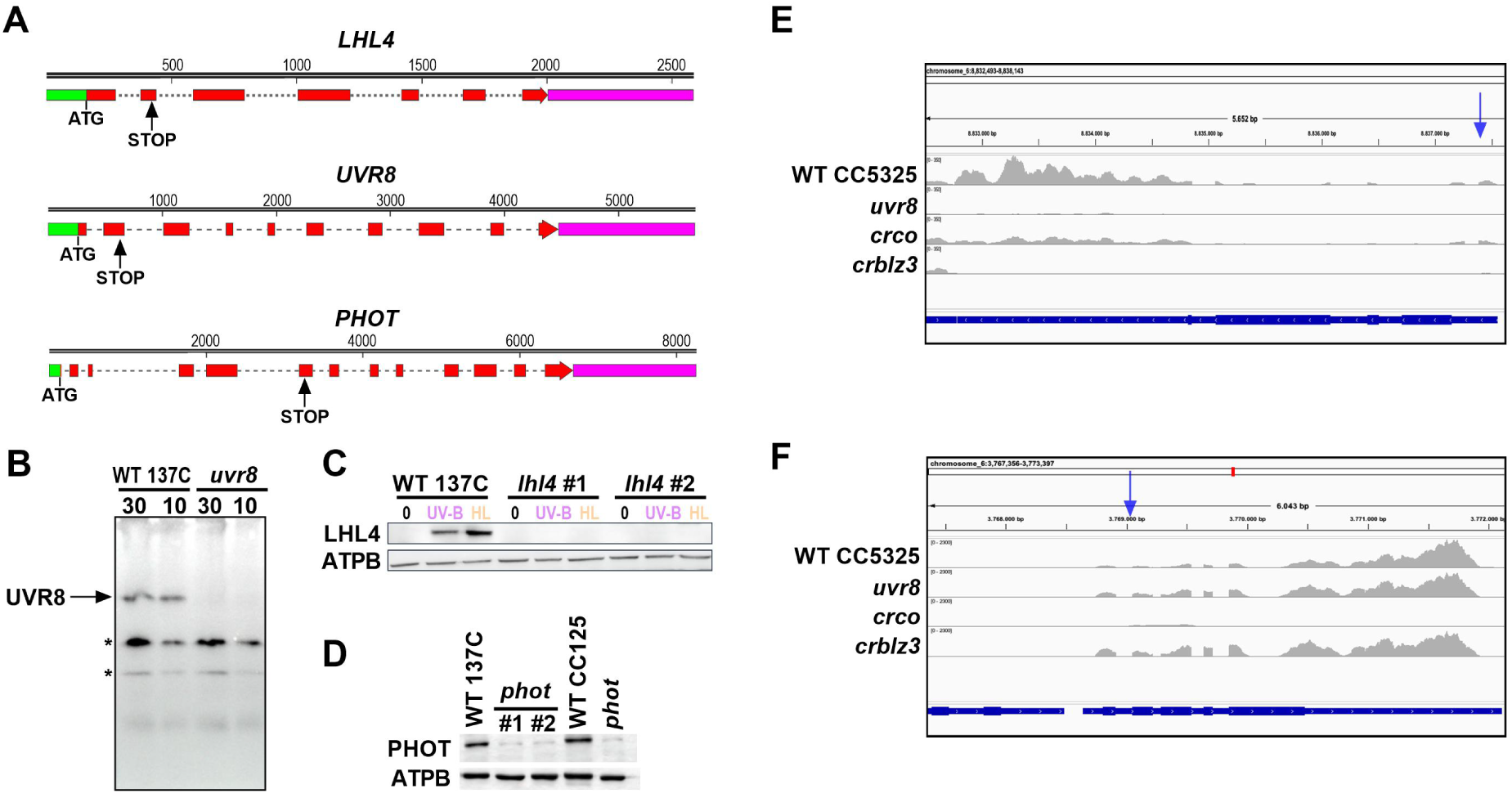
Validation of the mutants used and generated in this study. **(A)** Map of the *LHL4, UVR8* and *PHOT* genes. The positions of the translation initiation codon (ATG) of each gene, and stop codon inserted by CRISPR/Cas9 (STOP) are indicated. Green rectangle: 5’UTR; Pink rectangle: 3’UTR; Exons: red rectangles; Dotted lines: introns; numbers: bp. **(B)** Immunodetection of UVR8. Total proteins from WT and *uvr8* were extracted, and 10 or 30 µg were loaded onto the gel. The location of UVR8 is indicated. Asterisks indicate nonspecific bands. **(C)** Immunodetection of LHL4 in the WT and two independent *lhl4* mutants (#1 and #2). ATPB was used as loading control. **(D)** Immunodetection of PHOT. Total proteins of two independent *phot* mutants were loaded along with the previous mutant from Greiner et al., 2016 and their corresponding WT. The antibody recognizes a faint nonspecific band in all mutant strains. **(E-F)** Alignments of reads of WT, *uvr8, crco, crblz3* treated with UV-B at the **(E)** *Cre06.g310500* locus and **(F)** *Cre06.g278159* locus. Blue arrows indicate the position of the insertion for *crblz3* mutant (E) and *crco* mutant (F).

